# Monounsaturated fatty acid biosynthesis is critical for streptococcal envelope homeostasis and stress tolerance

**DOI:** 10.64898/2026.01.14.699484

**Authors:** Jonathon L. Baker, Jonah Tang, Mingzhe Guo, Felipe Fabrício Farias-da-Silva, Matthew Barbisan, Molly Burnside, Kaitlin Crofton, Samantha Williams, Saarin Rao, Michelle Lee, Sadie G. Drucker, Dustin Higashi, Justin Merritt, Yujiro Hirose, Jan-Willem Veening, Victor Nizet

## Abstract

The genus *Streptococcus* contains some of the most important commensals and pathogens of the human microbiome. To obtain the fatty acids required for cell membranes, *Streptococcus* either produce fatty acids *de novo* through the fatty acid biosynthesis (*fab*) pathway or uptake host fatty acids through the fatty acid kinase (*fak*) pathway. Although both the *fab* and *fak* pathways represent potential therapeutic targets to prevent or treat infection, progress is limited because of an incomplete understanding of taxon-to-taxon variability in streptococcal lipid metabolism. Here, we examined the role of *de novo* monounsaturated fatty acid (MUFA) synthesis in physiology and virulence-associated traits in *Streptococcus mutans, Streptococcus pyogenes,* and *Streptococcus pneumoniae,* three major pathogens that cause disease at distinct body sites. In all three species, deletion of *fabM* abolished MUFA production and caused severe growth defects, decreased stress tolerance, increased antibiotic susceptibility, and defects in cell viability, morphology, and division. In *S. mutans,* loss of *fabM* also markedly reduced competence signaling and production of the mutacin IV bacteriocin. Deletion of *fabM* increased susceptibility to killing by human neutrophils in *S. mutans* and *S. pneumoniae,* but not *S. pyogenes*. Together, these findings illustrate that MUFA synthesis is broadly important for streptococcal physiology and cell membrane homeostasis, while its contribution to pathogenesis is strongly species- and context-dependent, providing leads to guide development of novel therapeutic and/or preventative strategies.

## Introduction

The genus *Streptococcus* includes some of the most important pathogens and commensals of the human microbiota, inhabiting diverse niches and exerting profound effects on host health and disease. *Streptococcus pyogenes* (group A *Streptococcus*), *Streptococcus agalactiae* (group B *Streptococcus*), and *Streptococcus pneumoniae* all appear on both the 2024 WHO Priority Pathogens List and the CDC Antibiotic Resistance Threats Report. To persist in their ecological niches and, in the case of pathogenic species, cause disease, streptococci must withstand a range of environmental stresses. As Gram positive bacteria, *Streptococcus* spp. are surrounded by a cell envelope consisting of a single cytoplasmic membrane (i.e., lipid bilayer) and a thick peptidoglycan cell wall; many species *additionally* produce a capsule that is critical for virulence.

The lipid bilayer of the cytoplasmic membrane is composed of diverse lipid species, typically consisting of one to four fatty acid chains attached to a head group (T.-H. Lee et al. 2024). Fatty acid chains may vary in length and saturation. Saturated fatty acids (SFAs) adopt relatively straight configurations that allow tight packing and reduce membrane fluidity, whereas. unsaturated fatty acids contain one or more double bonds that may introduce bends into the hydrocarbon chain, increasing steric hinderance and membrane fluidity (T.-H. Lee et al. 2024). In *Streptococcus* spp., fatty acids can either be made *de novo* via the fatty acid biosynthesis (*fab*) operon (Parsons and Rock 2013) or incorporated from the environment through the fatty acid kinase (*fak*) system (Gullett et al. 2019; Radka 2023; Shi et al. 2022). De novo synthesis in streptococci is limited to SFAs and monounsaturated fatty acids (MUFAs). These fatty acids are attached to lipid head groups which, in *Streptococcus* spp., include glycerophosphates, sugars, and amino acids (Custer et al. 2014), through the *pls* machinery (Lu et al. 2006). By mixing and matching different fatty acid chains and head groups, bacteria can modulate membrane fluidity, curvature, and charge (T.-H. Lee et al. 2024), thereby promoting survival under environmental stresses such as temperature extremes, acid stress, reactive oxygen species, and antibiotic exposure.

Previous research demonstrated that *Streptococcus gordonii, S. salivarius, S. mutans* and *Lactobacaillus casei* increase the proportion of MUFAs in their membranes in response to environmental acidification, largely at the expense of SFAs (Fozo, Kajfasz, and Quivey 2004). In *S. mutans,* a similar shift toward increased MUFA content occurs under oxidative stress (Quivey et al. 2015; Derr et al. 2012). MUFAs are synthesized *de novo* in streptococci via the FabM enolyl-ACP isomerase, which was initially characterized biochemically using purified protein encoded by the *S. pneumoniae* gene sequence (Marrakchi, Choi, and Rock 2002) and later through genetic disruption in *S. mutans* (Fozo and Quivey 2004). In *S. mutans,* deletion of *fabM* resulted in membranes composed entirely of SFAs, accompanied by impaired tolerance to acid and oxidative stress and markedly reduced virulence in a rat model of dental caries (Fozo and Quivey 2004; Fozo et al. 2007). In contrast, deletion of *fabM* (as well as Δ*fabF* and Δ*fabMF*) in *S. agalactiae* did not attenuate virulence in a neonatal mouse sepsis model (Brinster et al. 2009). This discrepancy was hypothesized to reflect differences in concentration and composition of environmental fatty acids available at distinct infection sites, as fatty acid levels in saliva and dental plaque are substantially lower than those in blood (Brinster et al. 2009).

These observations raised important questions regarding the suitability of fatty acid biosynthesis enzymes as therapeutic targets. The *fab* enzymes differ substantially from their eukaryotic counterparts, making them attractive candidates for selective inhibition (Brinster et al. 2009; Parsons and Rock 2011; Radka and Rock 2022). However, the ability of some streptococci to bypass de novo fatty acid synthesis by scavenging host-derived fatty acids through the *fak* system has complicated this paradigm. Subsequent studies have highlighted that the efficacy of Fab inhibitors (e.g., platensimycin) is highly dependent on both bacterial species and infection context, including the availability of exogenous fatty acids at the infection site (Morvan et al. 2016; Lambert et al. 2024; Parsons et al. 2011; Radka and Rock 2022).

In this study, we further investigated the role of *de novo* MUFA biosynthesis in streptococcal physiology and virulence by examining *fabM* deletion mutants in *S. mutans, S. pyogenes,* and *S. pneumoniae,* three pathogens adapted to distinct anatomic niches. While *S. mutans* is a primary etiologic agent of dental caries and occasionally endocarditis (Lemos et al. 2019), *S. pyogenes* causes diseases ranging from pharyngitis and impetigo to necrotizing fasciitis and toxic shock syndrome (Brouwer et al. 2023), and *S. pneumoniae* is a leading cause of pneumonia, otitis media, and meningitis (Narciso et al. 2025). We show that, similar to *S. mutans*, deletion of *fabM* in *S. pyogenes* and *S. pneumoniae* abolishes MUFA production and severely impairs growth, phenotypes that could be rescued to varying degrees by exogenous unsaturated fatty acids. Across all three species, *fabM* deletion increases susceptibility to environmental stresses and antibiotics and is associated with defects in cell division, morphology, and viability. However, the ability of some streptococci to bypass de novo fatty acid synthesis by scavenging host-derived fatty acids through the *fak* system has complicated this paradigm.

## Results

Deletion of *fabM* in *S. mutans, S. pyogenes,* and *S. pneumoniae* eliminates production of UFAs and significantly impairs growth. A *fabM* deletion mutation in *S. mutans* (*SmΔfabM*) was constructed as part of the single-gene knockout library in *S. mutans* UA159 described in (Quivey et al. 2015). This mutant was significantly impaired in overall growth, defective in biofilm formation, and markedly less tolerant to acid and oxidative stress (Quivey et al. 2015). Consistent with prior reports using an insertional *fabM*::erm mutant (Fozo and Quivey 2004; Baker, Abranches, et al. 2015), deletion of *fabM* abolished production of UFAs in *S. mutans* (Figure 1).

**Figure 1:**
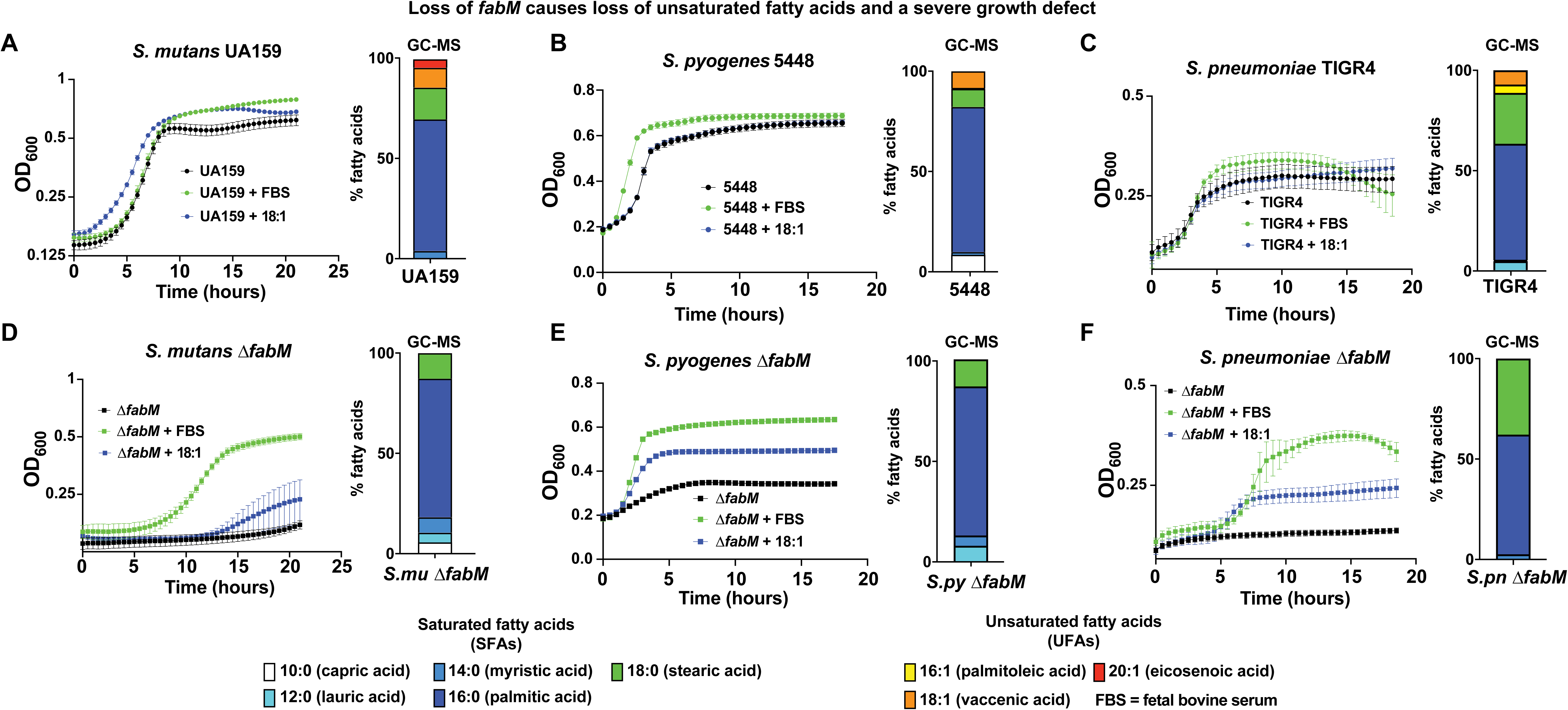
Δ*fabM* strains have impaired growth and do not produce MUFAs. Growth curves of the indicated parental strains (Panels A-C) or Δ*fabM* mutant strains (Panels D-F) in DMEM with and without supplementation of either 10% FBS fetal bovine serum (FBS) or 10 µg/mL vaccenic acid (18:1cis11). Bar graph insets to the right of each growth curve display in the relative abundances of fatty acids present in the cognate strain at the end of the growth curve in the unsupplemented DMEM as determined by GCMS. The legend for these bar charts is at the bottom of the figure.

To assess whether *de novo* MUFA biosynthesis plays a similar role in streptococcal pathogens adapted to cause infection at other body sites—and therefore exposed to distinct types and concentrations of environmental fatty acids, single-gene *fabM* deletion mutants were also created in the highly virulent M1T1 serotype *S. pyogenes* strain 5448 (*SpyΔfabM*;) and capsule serotype 4 *S. pneumoniae* strain TIGR4 (*SpnΔfabM*). As observed for *SmΔfabM,* both *SpyΔfabM* and *SpnΔfabM* failed to produce UFAs and exhibited significant growth defects in unsupplemented media (Figure 1).

In all three species, the growth defect of the Δ*fabM* strains could be rescued nearly to parental levels by supplementation of the growth media with fetal bovine serum (FBS), which contains both MUFAs and PUFAs. Growth was also improved by addition of purified vaccenic acid (18:1cis11), a UFA that is predominantly synthesized de novo by streptococci (Figure 1). A complemented *Smu*Δ*fabM* strain has been described previously and restored MUFA production, growth, and acid and oxidative stress tolerance to a large degree (Fozo and Quivey 2004). A complemented strain of *Spn*Δ*fabM* constructed in this study similarly restored MUFA production and largely suppressed the impaired growth phenotype (Figure S1). A complemented strain of *SpyΔfabM* was not generated due to the technical challenges associated with genetic manipulation of strain 5448; however, the concordant phenotypic rescue observed in *S. mutans* and *S. pneumoniae* supports that the observed phenotypes are attributable specifically to loss of *fabM*. The ability of other unsaturated fatty acids to rescue growth of the Δ*fabM* strains was also examined (Figure 2). In all three species, the MUFAs palmitoleic acid (16:1cis9), oleic acid (18:1cis9), and eicosenoic acid (20:1cis11) rescued growth in a dose-dependent manner, with maximal rescue observed at concentrations up to 40 µg/ml for *Smu*Δ*fabM* and *Spn*Δ*fabM,* and up to 10 µg/ml for *Spy*Δ*fabM* (Figure 2). The diene PUFA linoleic acid (18:2cis 9,12), which is abundant in human blood, also rescued growth, but over a narrower concentration range. Linoleic acid rescued growth of *Smu*Δ*fabM* in a dose-dependent manner up to 20 µg/ml, whereas higher concentrations were inhibitory. In contrast, growth of *Spy*Δ*fabM* and *Spn*Δ*fabM* was rescued only at low concentrations (≤ 2.5 µg/ml) with higher concentrations strongly inhibitory (Figure 2).

**Figure 2:**
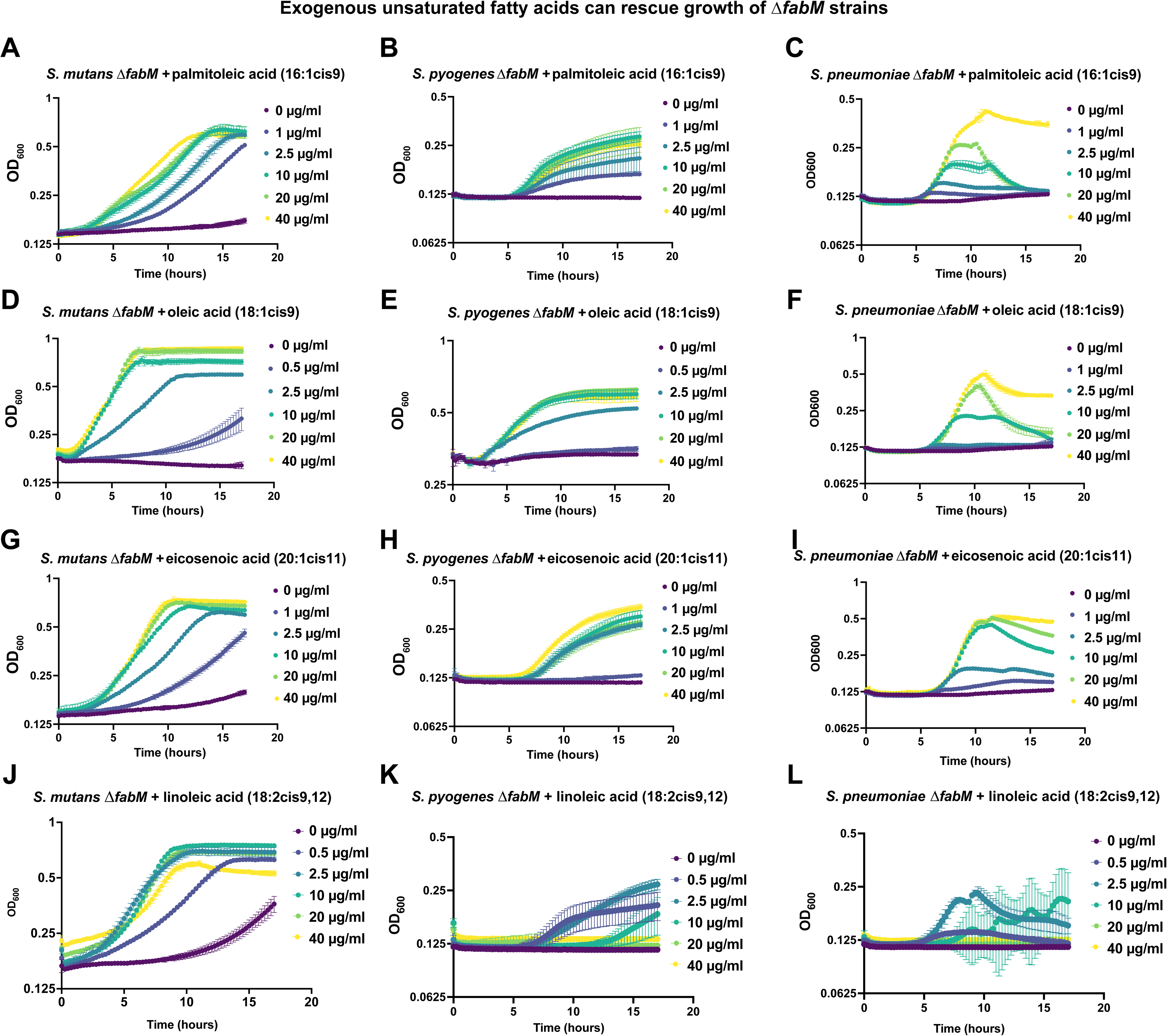
Exogenous monounsaturated and polyunsaturated fatty acids can rescue growth of Δ*fabM* strains. Growth curves of the indicated strains with the indicated supplements at the indicated concentrations.

Streptococci are not known to synthesize PUFAs *de novo* but can incorporate exogenous PUFAs through the *fak* system. Notably, linoleic acid rescued growth of *Smu*Δ*fab,* despite the absence of *fakB3,* the gene encoding the PUFA-incorporating FakB component characterized in *S. pneumoniae* (Gullett et al. 2019). This finding suggests that *S. mutans* may encode an alternative mechanism for PUFA incorporation. Although *S. pyogenes* encodes a *fakB3* homolog, the role of this gene in PUFA utilization for this species has not been directly examined. As expected, provision of the SFA stearic acid (18:0) did not rescue growth of any Δ*fabM* strain (data not shown).

### Deletion of *fabM* in *S. mutans, S. pyogenes,* and *S. pneumoniae* increases susceptibility to acid stress, oxidative stress, and antibiotics

An insertional mutation of *fabM* in *S. mutans* was previously shown to impair tolerance to environmental stresses such as acid and oxidative stress (Fozo and Quivey 2004). To determine whether deletion of *fabM* produces similar phenotypes across streptococcal species, the stress tolerance of *ΔfabM* strains and their cognate parent strains was examined under conditions of temperature stress, acid stress, oxidative stress, and antibiotics exposure.

In *S. mutans,* SmuΔ*fabM* exhibited no detectable growth at 30°C, while growth at 42°C was improved relative to growth at 37°C (Figure 3). Growth of *Smu*Δ*fabM* was also reduced to nearly undetectable levels at pH 5.4 and in the presence of 1 mM H_2_O_2_, consistent with previous reports describing the *Smu*Δ*fabM* phenotype. In contrast, *SpyΔfabM* and *SpnΔfabM* exhibited only slightly reduced growth at 30°C relative to 37°C and showed little to no growth at 42°C (Figure 3). All three Δ*fabM* strains exhibited pronounced sensitivity to acid stress, with little to no growth at pH 6.0 (Figure 3).

**Figure 3:**
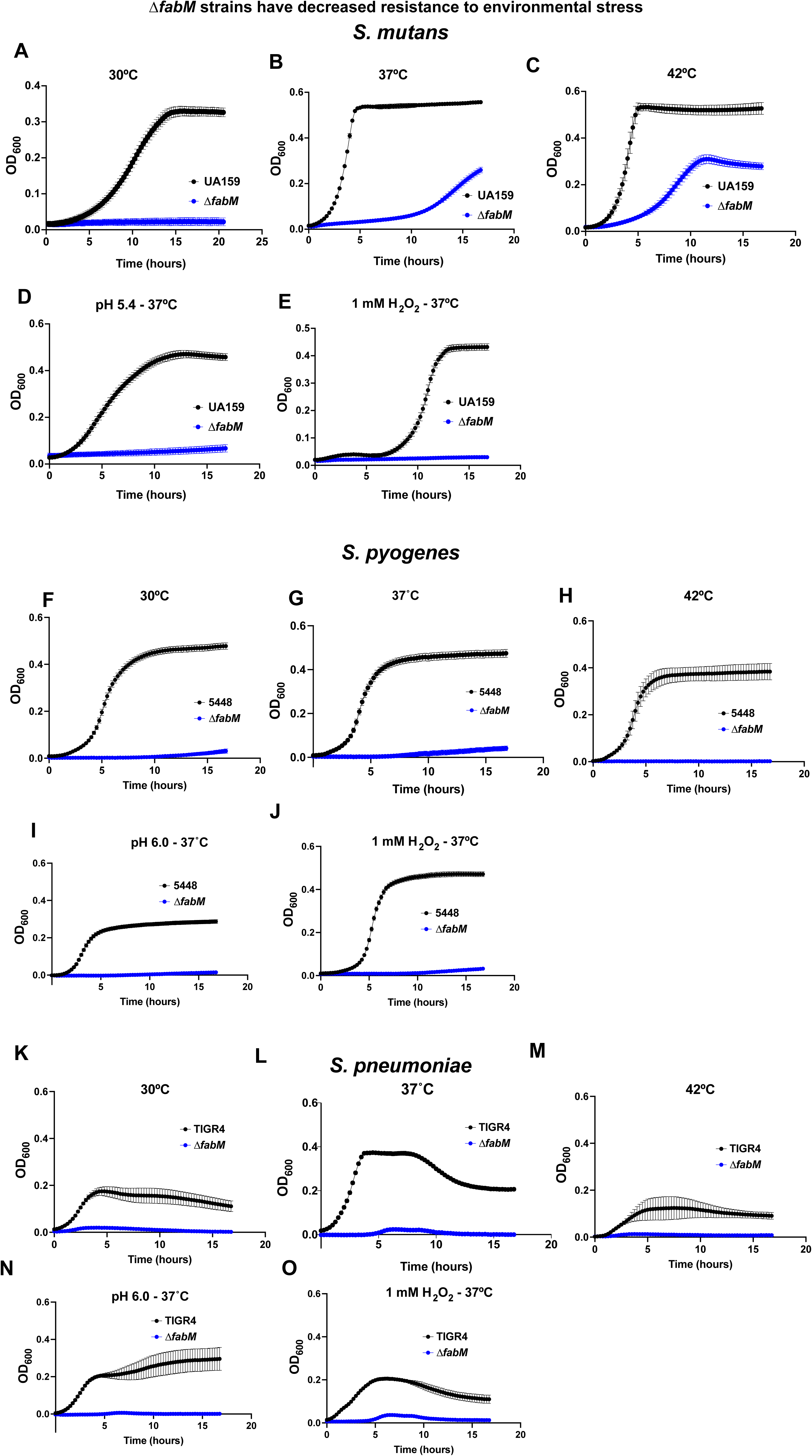
Δ*fabM* strains are sensitive to adverse temperature, acid, and oxidative stress. Growth curves of the indicated strains of *S. mutans* (Panels A-E), *S. pyogenes* (Panels F-J), or *S. pneumoniae* (Panels K-O) grown in THB at the indicated temperatures (Panels A,B,C,F,G,H,K,L,M), grown at 37°C in THB buffered to the indicated acidic pH (Panels D,I,N), or grown at 37°C in THB + 1 mM H_2_O_2_ (Panels E,J,O).

*SpyΔfabM* and *SpnΔfabM* were less sensitive to H_2_O_2_ than *SmuΔfabM.* This difference is likely attributable to that fact that *S. pyogenes* and *S. pneumoniae* can produce much higher levels of H_2_O_2_ than *S. mutans and* therefore encode several mechanisms to tolerate oxidative stress. To assess whether loss of MUFA synthesis also alters antibiotic susceptibility, minimum inhibitory concentrations (MICs) were determined for antibiotics representing multiple mechanistic classes: ampicillin, daptomycin, polymyxin B, and spectinomycin. In all three species MICs for daptomycin, polymixin B, and spectinomycin were reduced in the Δ*fabM* strains compared to their cognate parent strains (Table 1). For ampicillin, MICs were reduced in the Δ*fabM* strains of *S. mutans* and *S. pyogenes*, whereas for *S. pneumoniae* both TIGR4 and *SpnΔfabM* exhibited MICs below the limit of detection (Table 1).

**Table 1:**
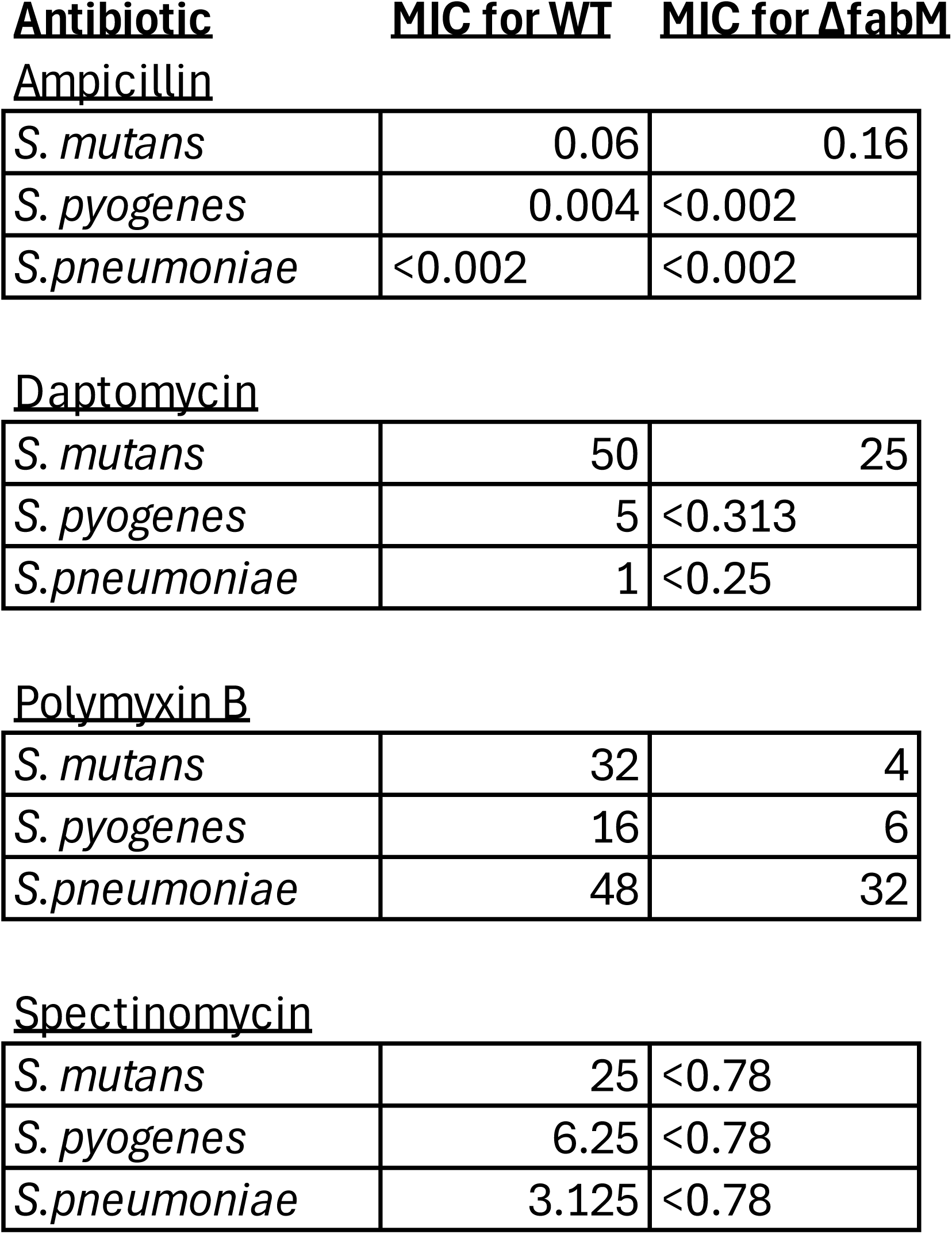
MIC of indicated strains for indicated antibiotic **<u>Antibiotic</u> <u>MIC</u>** <u>**for WT**</u> **<u>MIC</u>** <u>**for &fabM**</u> <u>Ampicillin</u>

|  |  |  |
| --- | --- | --- |
| *S. mutans* | 0.06 | 0.16 |
| *S. pyogenes* | 0.004 | <0.002 |
| *S.pneumoniae* | <0.002 | <0.002 |
| Daptomycin |  |  |
| *S. mutans* | 50 | 25 |
| *S. pyogenes* | 5 | <0.313 |
| *S.pneumoniae* | 1 | <0.25 |
| Polymyxin B |  |  |
| *S. mutans* | 32 | 4 |
| *S. pyogenes* | 16 | 6 |
| *S.pneumoniae* | 48 | 32 |
| Spectinomycin |  |  |
| *S. mutans* | 25 | <0.78 |
| *S. pyogenes* | 6.25 | <0.78 |
| *S.pneumoniae* | 3.125 | <0.78 |

Taken together, these data indicate that loss of *fabM* and the absence of *de novo* MUFA production broadly sensitize streptococcal pathogens to environmental stresses and multiple classes of antibiotics, consistent with compromised membrane and cell envelope homeostasis.

### Mutacin production and competitive fitness versus neighboring commensals is impaired in *Smu*Δ*fabM*

Many *Streptococcus* species produce bacteriocins that inhibit the growth of neighboring bacteria, often other streptococci *inhabiting the same ecological niche*. To assess whether deletion of *fabM* impacts competitive fitness, *Smu*Δ*fabM* was examined in a standard competition assay on THB agar commonly used to measure inter-streptococcal antagonism. *Smu*Δ*fabM* was significantly less effective than the parent UA159 strain at inhibiting growth of the oral commensals *Streptococcus gordonii, S. sanguinis,* and *S. mitis* (shown in Figure 4A for *S. sanguinis*; data not shown for *S. gordonii* and *S. mitis*). Although supplementation of the growth medium with MUFAs and/or PUFAs partially improved competitive inhibition by *Smu*Δ*fabM*, these effects did not reach statistical significance due to assay variability (Figure 4A). Interestingly, a similar increase in inhibitory activity was observed for the UA159 parent strain upon fatty acid supplementation (Figure 4A).

**Figure 4:**
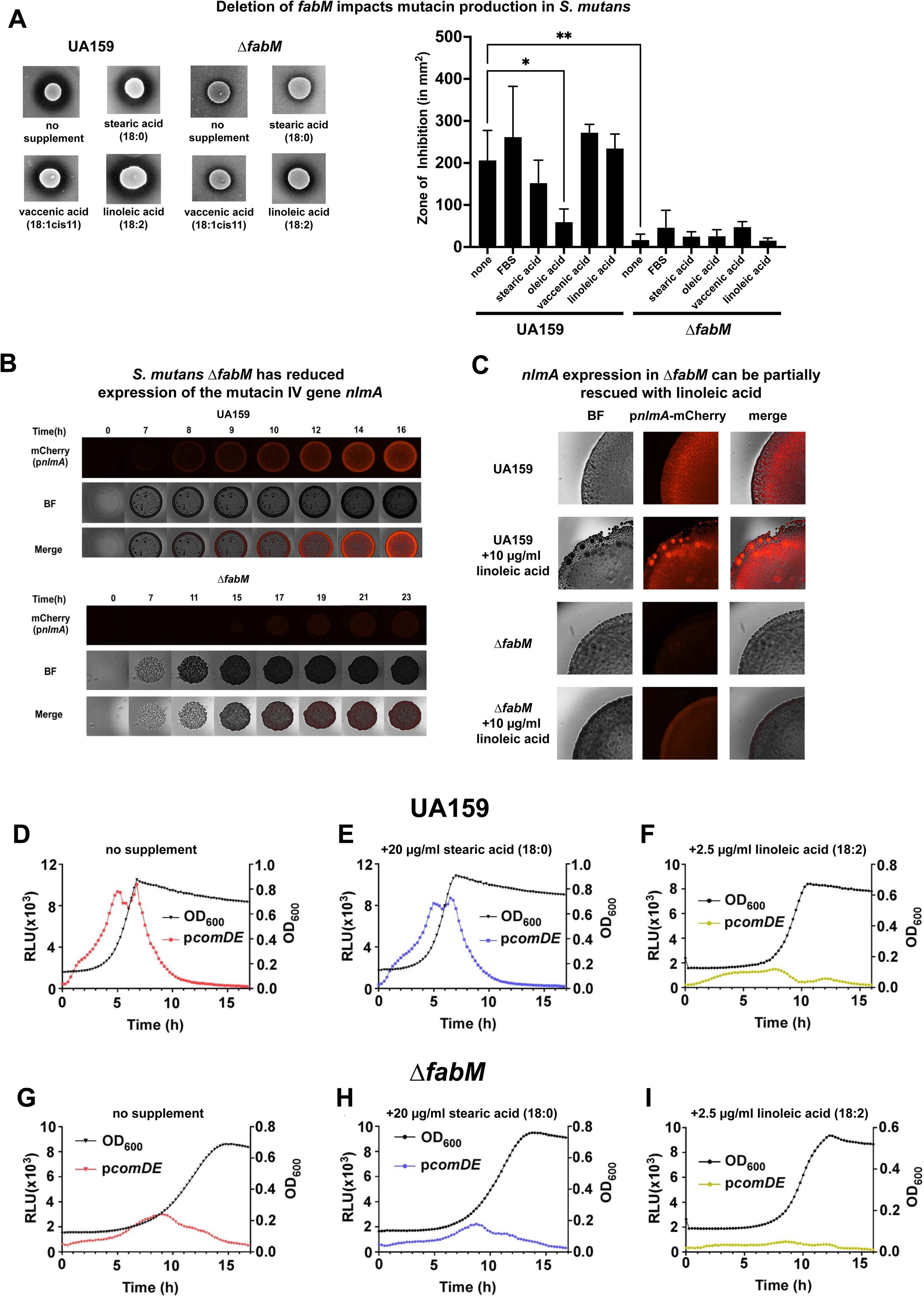
Deletion of *fabM* in *S. mutans* impacts mutacin production and competence signaling. Panel A includes representative photos of the zone of inhibition produced by 8 µl colonies of indicated strain spotted on THB agar with the indicated supplements with a *S. sanguinis* overlay in soft agar. Bar graph on the right side of Panel A is quantification of 3 experimental replicates of the indicated strain/conditions of the same competition assay between *S. mutans* and *S. sanguinis* displayed in the Panel A photos. Panel B shows representative photos of a time course of growth of a p*nlmA-*mCherry reporter in either a UA159 or *Smu*Δ*fabM,* as indicated. BF, bright field. Panel C is representative images of the p*nlmA-*mCherry reporter in the indicated background strain, with or without supplementation of 10 µg/ml linoleic acid, as indicated. Panels D-I are superimposed growth curves and luminescence produced by a p*comDE-*luciferase reporter, with the indicated supplements, in either a UA159 (Panels D-F) or Δ*fabM* (Panels G-I) background.

Because *S. mutans* mutacin IV is the primary bacteriocin responsible to inhibition of the tested commensal *Streptococcus* spp., we examined the expression of the *nlmA* component of mutacin IV using a P*_nlmA_-mCherry* transcriptional reporter. In the UA159 background, robust mCherry fluorescence was readily observed overlaid with colonies grown on agar plates (Figure 4B). In contrast, *Smu*Δ*fabM* exhibited negligible mCherry fluorescence, indicating a marked reduction in *nlmA* expression. Expression of mCherry in the Δ*fabM* background could be rescued to a modest degree by provision of linoleic acid (Figure 4B and 4C).

In *S. mutans,* mutacin production is typically controlled by the competence-associated two-component system ComDE. To determine whether deletion of *fabM* affects ComDE signaling, P*_comDE_-*luciferase reporter strains were generated in both the UA159 and *Smu*Δ*fabM* backgrounds. Expression of *comDE* was significantly reduced in *Smu*Δ*fabM* grown in unsupplemented media as well as media supplemented with stearic acid (Figure 4D, 4G, 4E, and 4H), consistent with the observed reduction in *nlmA* expression and mutacin production. Unexpectedly, addition of linoleic acid reduced, rather than increased, *comDE* expression in both the UA159 and Δ*fabM* backgrounds (Figure 4F and 4I). This contrasted with the partial restoration of *nlmA* expression in the Δ*fabM* background upon linoleic acid supplementation, suggesting that linoleic acid may decouple *nlmA* expression from canonical ComDE-dependent regulation.

The *S. pyogenes* and *S. pneumoniae* strains examined in this study did not inhibit growth of the tested commensal streptococci under the assay conditions used (data not shown). Because *S. pneumoniae* and *S. pyogenes* typically cause α and β hemolysis, respectively, the impact of *fabM* deletion on hemolysis was also examined using THB agar supplemented with blood. In both cases hemolysis was not affected, potentially reflecting complementation of the *ΔfabM* phenotype by UFAs present in blood (data not shown).

### Δ*fabM* strains exhibit disrupted cehll morphology and altered CFU/OD_600_ ratios

Initial examination of the Δ*fabM* strains by light microscopy revealed abnormal cell morphology in all three species. Although chains of cells formed by the *S. mutans* and *S. pyogenes* Δ*fabM* strains appeared shorter than those of their cognate parent strains, this difference was not statistically significant due to substantial variability in chain length (Figure 5A). Supplementation of the growth medium with UFAs (vaccenic acid or FBS) resulted in significantly longer chains in *S. pyogenes* 5448, but not in the *SpyΔfabM* strain (Figure 5A). Deletion of *fabM* did not measurably impact chain length in *S. pneumoniae,* which typically forms very short chains under the conditions tested (data not shown).

**Figure 5:**
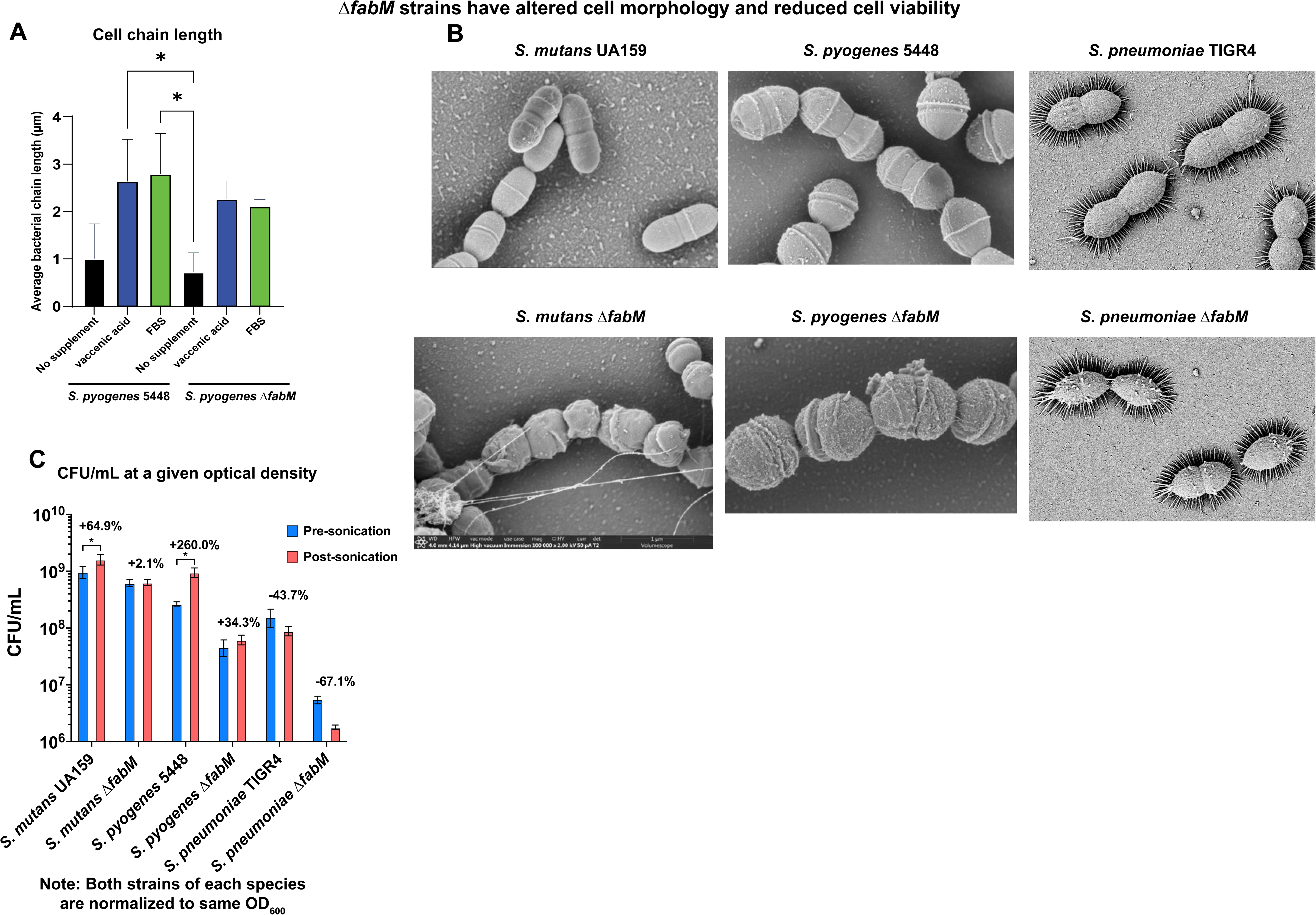
Deletion of *fabM* impacts cell morphology. Panel A is a bar graph illustrating the effect of *fabM* deletion and/or UFA supplementation on cell chain length in *S. pyogenes.* *denotes statistical significance (p < 0.05) between indicated groups of samples based on a Tukey’s Honestly Significant Differences test following a one-way ANOVA. Panel B is representative images of scanning electron microscopy of the indicated strains. Note asymmetry of cells and divisomes in the Δ*fabM* strains, as well as additional membrane blebs in the *SpnΔfabM* strain, specifically. “Spikes” surrounding the *S. pneumoniae* cells are believed to be an artefact of the capsule and fixation process. Images were taken at 80,000X. Panel C is a bar graph of the CFU/ml of the indicated strain, either pre- or post-sonication, as indicated. Both strains of each species are normalized to the same OD_600_. *denotes statistical significance (p < 0.05) between indicated groups of samples based on a Tukey’s Honestly Significant Differences test following a one-way ANOVA.

To examine cellular morphology at higher resolution, scanning electron microscopy was performed. In all three species, Δ*fabM* strains displayed pronounced morphological abnormalities compared to their respective parent strains (Figure 5B). In *Smu*Δ*fabM* and *Spy*Δ*fabM*, cells frequently exhibited asymmetrical septa and in some cases appeared to initiate division without completing septation, suggestive of impaired cell division (Figure 5B). In *Spn*Δ*fabM,* cells appeared shorter and more ovoid and exhibited significantly more membrane blebs relative to TIGR4 (Figure 5B). Both *S. pneumoniae* strains exhibited spike-like structures surrounding the cells, which are believed to represent fixation artefacts associated with the capsule (Kuipers et al. 2016).

During routine normalization of cultures based on optical density (OD_600_), an inconsistency became apparent between OD_600_ and viable cell counts in the Δ*fabM* strains. Quantification revealed a significant reduction in the ratio CFU/OD_600_ ratio for all three Δ*fabM* strains compared to their cognate parent strains: a 1.67-fold decrease for *Smu*Δ*fabM*, a 3.86-fold decrease for *Spy*Δ*fabM*and a 23.63-fold decrease for *Spn*Δ*fabM* (Figure 5C). These findings indicate that, for a given culture density, a substantially smaller proportion of cells in the Δ*fabM* cultures are capable of forming colonies.

Because *Streptococcus* spp. often form chains, sonication is routinely used prior to CFU enumeration to break up chains and improve quantification. After sonication, CFU counts increased dramatically for the UA159 and 5448 parent strains, whereas only modest increases were observed for their corresponding Δ*fabM* strains (Figure 5C). This difference is consistent with the shorter average chain length observed in the Δ*fabM* strains and/or increased sensitivity of these strains to sonication. In contrast, sonication reduced CFU counts for both TIGR and *Spn*Δ*fabM*, indicating that the sonication procedure employed was detrimental to *S. pneumoniae* viability, with the *Spn*Δ*fabM* strain exhibiting greater sensitivity (Figure 5C).

Taken together, these results indicate that *de novo* MUFA synthesis is required for proper cell morphology and division in streptococci. Loss of *fabM* results in defective septation, abnormal cell shape, and a marked reduction in the fraction of viable cells within the population, consistent with compromised membrane function and cell envelope integrity.

### *Smu*Δ*fabM* and *Spn*Δ*fabM,* but not *SpyΔfabM* are more susceptible to neutrophil killing

Because loss of MUFA synthesis is expected to change properties of the cell envelope, which may in turn influence susceptibility to host immune defenses, the Δ*fabM* strains and their cognate parent strains were examined in a neutrophil killing assay using primary human neutrophils. *Smu*Δ*fabM* and *Spn*Δ*fabM* were readily killed by neutrophils, whereas the corresponding parent strains were not susceptible and exhibited CFU counts comparable to control conditions lacking neutrophils (Figure 6A). In contrast, neither *S. pyogenes* 5448 nor *Spy*Δ*fabM* exhibited detectable susceptibility to neutrophil-mediated killing under the conditions tested (Figure 6C).

**Figure 6:**
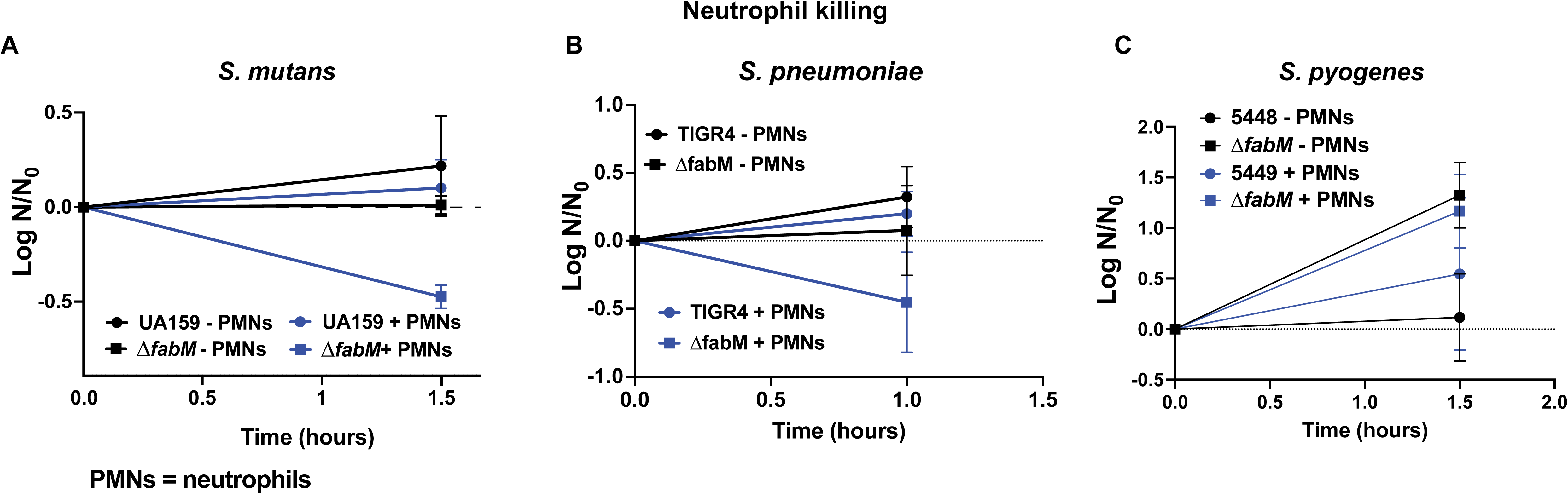
Deletion of *fabM* impacts susceptibility to neutrophil killing for *S. mutans* and *S. pneumoniae.* Panels A-C, graphs indicating survival of the indicated species and strain in either the absence or presence of neutrophils, as indicated.

## Discussion

As the physical barrier separating the inside and outside of the cell, the membrane plays a key role in physiology, and its properties have wide-reaching impacts on how cells interact with their surroundings (T.-H. Lee et al. 2024). Typical membrane lipids consist of fatty acid chains attached to a head group (T.-H. Lee et al. 2024). *Streptococcus* spp. are able to either generate SFA and MUFA chains *de novo* through the *fab* operon (Parsons and Rock 2013), or use the *fak* system to incorporate exogenous fatty acids from the environment (Gullett et al. 2019). The enoyl-ACP isomerase encoded by *fabM* is responsible for *de novo* biosynthesis of MUFAs, while *Streptococcus* are not known to produce PUFAs (Marrakchi, Choi, and Rock 2002). Several species of *Streptococcus* are known to increase the proportion of MUFAs in their membrane in response to environmental acidification. However, it remains unknown how this adjustment is controlled or precisely why this response is protective (Fozo, Kajfasz, and Quivey 2004; Pesakhov et al. 2007). In *S. mutans,* a *fabM* insertional mutant had no MUFAs in its membrane and exhibited impaired growth and acid tolerance, leading to reduced virulence in a rat model of dental caries (Fozo and Quivey 2004; Fozo et al. 2007). This was in stark contrast to a study that showed a *S. agalactiae* Δ*fabM* strain was fully virulent in a mouse neonatal sepsis model (Brinster et al. 2009). The same study also showed full virulence of Δ*fabF* and Δ*fabMF* strains, which would presumably be deficient in all *de novo* fatty acid synthesis, relying entirely on incorporation of exogenous fatty acids for growth (Brinster et al. 2009). The *fab* genes were initially touted as promising antibiotic targets due to their major structural differences from their eukaryotic counterparts (i.e., therefore inhibitors would likely have a low toxicity to mammalian cells) (Heath, White, and Rock 2001). However, the ability of certain species to circumvent the need for the *fab* genes by using exogenous fatty acids, as shown in the *S. agalactiae* study, brought the suitability of the *fab* genes as drug targets into question (Zlitni and Brown 2009). Subsequent research has illustrated that the utility of *fab* inhibitors is highly dependent on both the bacteria in question and the environmental context (Radka and Rock 2022; Parsons et al. 2013; Adams et al. 2021).

The goal of this study was to further characterize the role of MUFA production in *Streptococcus* physiology and virulence by continuing to determine the role of *fabM* in *S*. *mutans,* as well as explore the impact on Δ*fabM* deletion in *S. pyogenes* and *S. pneumoniae*. *S. mutans, S. pyogenes,* and *S. pneumoniae* have evolved to cause disease in different environments, each of which contain different types and concentrations of fatty acids. Similar to what was observed previously in *S. mutans* (Fozo and Quivey 2004), the Δ*fabM* strains in *S. pyogenes* and *S. pneumoniae* had severely impaired growth, which could be rescued to varying degrees by exogenous UFA supplementation. A previous study also examined *fabM* null strains in *S. mutans* and *S. pneumoniae* and reported that the strains were auxotrophs, with no growth in unsupplemented media (Altabe, Lopez, and de Mendoza 2007). The authors suggested that the differences observed between their study and the initial report of *SmuΔfabM,* which could grow (albeit not well) in unsupplemented media, could possibly be due to differences in the media recipe used — the Fozo & Quivey study using BHI and the Altabe et al. study using AGCHYE. That could also possibly explain differences seen between Altabe et al. and this study, as THB and DMEM were used here. It is also possible that there were phenotypic differences between the *S. mutans* UA159 isolates used in the Altabe et al. study and this study. Although the *SpnΔfabM* strain reported in this study barely grew in unsupplemented media, it was reported to be a complete auxotroph in Altabe et al (Altabe, Lopez, and de Mendoza 2007). Notably, the TIGR4 strain was used here, compared to that study, which used strain 709. The approach to mutant creation was also different, as the present study engineered an allelic replacement of the *fabM* ORF with an erythromycin resistance gene, while the Altabe et al. study created an insertional mutation to inactivate *fabM.* Again, the differences in these findings are likely explained by discrete isolate-to-isolate or environmental differences.

In terms of growth phenotypes, the *SmuΔfabM* strain exhibited no growth at 30°C and had a less severe growth phenotype at 42°C compared to 37°C. This makes sense, as a membrane composed entirely of SFAs would be more rigid (i.e. a higher melting point and less able to withstand colder temperatures). Indeed, other bacteria are known to increase the proportion of UFAs in their membranes in response to cold (Dong and Cronan 2021). With exclusively SFAs in *SmΔfabM,* it also makes sense that growing at higher-than-normal temperatures would be less of an issue. In contrast, *SpyΔfabM* and *SpnΔfabM* exhibited some growth at 30°C and no growth at 42°C. This could possibly be due to these strains evolving to reside in different environments with slightly different temperatures, such as the respiratory tract and skin. Notably, *S. mutans* in general had more MUFAs than both *S. pyogenes* and *S. pneumoniae*. *S. mutans* also produced 20-carbon fatty acids, mainly eicosenoic acid (20:1), along with trace amounts of eicosanoic acid (20:0). These 20-carbon fatty acids were not observed in *S. pyogenes* or *S. pneumoniae.* Similar to *S. mutans,* the *SpyΔfabM* and *SpnΔfabM* strains were impaired in their ability to withstand acid stress, exhibiting no growth at pH 6, although *S. mutans* was more acid tolerant overall, in line with previous observations (Lemos et al. 2019). Oxidative stress in the form of H_2_O_2_ significantly impaired growth of *SmuΔfabM,* but not *SpnΔfabM* or *SpyΔfabM*. While *S. mutans* does not produce high concentrations of H_2_O_2_, *S. pneumoniae*, and some strains of *S. pyogenes,* produce copious H_2_O_2_ through enzymes such as lactate oxidate, the peroxide-forming NADH oxidase, and pyruvate oxidase (*S. pneumoniae* only) (Henningham et al. 2015; Regev-Yochay et al. 2007; Taniai et al. 2008). *S. pyogenes* and *S. pneumoniae* may use H_2_O_2_ to inhibit the growth of competitors, and *S. pneumoniae* is known to use H_2_O_2_ to release and degrade heme, which is used as a source of iron (Alibayov et al. 2022; Womack et al. 2024). *S. pneumoniae* pyruvate oxidase is also related to capsule formation (Echlin et al. 2016) and pneumolysin production (Bryant et al. 2016). The H_2_O_2_ produced by *S. pneumoniae* appeared to modulate MUFA production as *S. pneumoniae* cells grown anaerobically, in acidic conditions, or with a mutation in pyruvate oxidase contained significantly more vaccenic acid (18:1cis11) in their membranes (Pesakhov et al. 2007). Furthermore, H_2_O_2_ was shown to oxidize a cysteine residue in FabF, altering the *S. pneumoniae* membrane composition (Benisty et al. 2010). This also aligns with *S. mutans* containing significantly more MUFAs when grown in conditions of oxidative stress (Derr et al. 2012; Baker, Derr, et al. 2015). Across all three species, Δ*fabM* strains had increased susceptibility to antibiotics. Initially, it was expected that this might be limited to antibiotics targeting the cell envelope structures (e.g., membrane, or cell wall), such as ß-lactams, polymyxin B, or daptomycin, especially since membrane composition is known to impact some antibiotic susceptibilities, especially for daptomycin (Hines et al. 2017). However, Δ*fabM* appeared to exhibit a general increase in susceptibility to all antibiotics (and stress in general) as all three Δ*fabM* strains had a lower MIC for the bacterial ribosome inhibitor spectinomycin.

The effect of *fabM* deletion on the ability of *S. mutans, S. pneumoniae,* and *S. pyogenes* to inhibit the growth of competing oral commensal *Streptococcus* spp. was also examined. Although *S. pneumoniae* and *S. pyogenes* did not inhibit the tested commensal oral *Streptococcus* in our assays, *S. mutans* produces bacteriocins that readily inhibit the growth of these organisms (Lemos et al. 2019). *SmuΔfabM* was significantly impaired in its ability to inhibit the growth of the commensal *Streptococcus*. As mutacin production in *S. mutans* is frequently controlled by the two-component ComDE quorum sensing system (Lemos et al. 2019; Merritt and Qi 2012), the impact of Δ*fabM* on *comDE* expression was examined. *comDE* expression was significantly reduced in the Δ*fabM* strain, which may explain the reduction in *nlmA* expression and mutacin IV production. However, addition of linoleic acid also reduced *comDE* expression, indicating that the rescue of *nlmA* expression observed upon addition of linoleic acid must have been independent of *comDE* signaling. The observation that linoleic acid inhibited the *comDE* system was especially interesting, as this system is highly involved in virulence traits such as biofilm formation, competence, and toxin production (Merritt and Qi 2012). Although unsaturated fatty acids have been shown to impact quorum sensing in *Pseudomonas aeruginosa* (Han et al. 2024) and *Acinetobacter baumanii* (Nicol et al. 2018), there have been no such reports in *Streptococcus*. However it is notable that short chain fatty acids impact *comDE* in *S. gordonii* (Park et al. 2021) and that the BriC quorum sensing peptide regulates fatty acid biosynthesis in *S. pneumoniae* (Aggarwal et al. 2021). The impact of long chain SFAs and UFAs on *Streptococcus* quorum sensing warrants further study which is currently in progress.

Routine examination of the Δ*fabM* strains under a light microscope appeared to show abnormal cell morphology and reduced cell chain lengths. Quantification of cell chain lengths indicated that the reduction observed in the Δ*fabM* strains was not statistically significant, likely due to high variability of chain length. Scanning electron microscopy was utilized to observe the cells at a higher resolution. While the cells of the parent strains had regular shapes with symmetrical septa and divisomes, most Δ*fabM* cells were asymmetrical — some with multiple septa, especially in *SmuΔfabM* and *SpyΔfabM*. In *SpnΔfabM,* cells appeared to have a shorter, more ovoid shape and had a greater number of membrane blebs compared to TIGR4. Collectively, these images indicate that the Δ*fabM* strains have a defect in cell division and/or morphology. In *S. pneumoniae,* it was reported that SFAs tended to co-localize with the cell-division machinery, while UFAs made up the other regions of the cell (Calvez et al. 2019). It seems likely that in the absence of UFAs, membrane fluidity is abnormal, and the cell division machinery has difficulty localizing or operating properly. This seemed to translate to an abundance of cells in the cultures that were apparently not viable. Furthermore, when cultures of the cognate parent strains and Δ*fabM* strains were normalized to the same culture density, the number of CFU/ml of the Δ*fabM* strains was significantly lower, indicating that many of the cells in the Δ*fabM* strains are not viable. Sonication of the cells to break up the chains is commonly used prior to quantification of *Streptococcus* spp. by CFU plating. This sonication increased the number of CFU in the parent strains of *S. mutans* and *S. pyogenes* but did so to a lesser extent in the Δ*fabM* strains, possibly due to the shorter chain length in the Δ*fabM* strains. Alternatively, the Δ*fabM* cells could be more sensitive to sonication, and the process may kill some of the cells. Notably, *S. pneumoniae* decreased in CFU/ml following sonication, indicating it may be more sensitive to the sonication protocol.

The *S. mutans* and *S. pneumoniae ΔfabM* strains were more susceptible to killing by human neutrophils, likely reflective of the general susceptibility to stress observed in the Δ*fabM* mutants. It is unclear exactly why the *S. pyogenes ΔfabM* strain was still resistant to neutrophil killing, however *S. pyogenes* is known to employ several mechanisms to avoid this fate including evasion of phagocytosis, resistance to antimicrobial peptides, and escape from neutrophil extracellular traps (Bergsten and Nizet 2024; Henningham et al. 2015).

Research further examining the ability of *S. mutans* to utilize PUFAs while lacking a *fakB3* gene, as well as further study of the impact of exogenous fatty acids on *Streptococcus* competence signaling and virulence are currently in progress. A limitation of this study was that while GCMS analysis of fatty acid methyl esters (the technique employed here) captures all fatty acid tails in the sample, it ignores head groups of the lipid molecules, which undoubtedly also impact cellular physiology. However, there is presently no single chromatography-mass spectrometry method that captures all lipids, as those lipids with different classes of head groups need to be analyzed separately with current technology. Therefore, to capture the most complete set of data about the fatty acid chains themselves, GCMS was the technique of choice here. Overall, this study also suggested that UFAs play an important role in the organization and function of the cell division machinery. As data accumulates showing that the effectiveness of fatty acid biosynthesis inhibitors is highly dependent on species and disease context, examining further relevant situations such as lung infection by *S. pneumoniae* or skin infection by *S. pyogenes* is likely to open the door to novel context- and pathogen-specific preventative and therapeutic strategies.

## Materials and Methods

### Bacterial strains and growth conditions

All strains, plasmids, and primers used in this study are listed in Supplemental Table S1. All *Streptococcus* strains were maintained on Todd-Hewitt broth (THB) agar plates (BD/Difco, Franklin Lakes, NJ) at 37°C in a 5% (vol/vol) CO_2_–95% air environment. The *S. mutans* strains UA159 (Ajdić et al. 2002) and *Smu*Δ*fabM* (Fozo and Quivey 2004; Quivey et al. 2015) have been described previously. An *SmuΔfabM* complement strain, which faithfully restored production of MUFAs and most other phenotypes of the *fabM* deletion has also been previously described (Fozo and Quivey 2004).

### Generation of recombinant strains: *S. pyogenes* Δ*fabM*

Genomic DNA from *S. pyogenes* 5448 (Chatellier et al. 2000) was isolated using the Quick-DNA Fungal/Bacterial Miniprep Kit (Zymo, Inc.) according to the manufacturer’s instructions. 500 bp upstream of *fabM* was amplified using PCR primer pair Spy-fabM-LF-F and Spy-fabM-LF-R, 500 bp downstream of *fabM w*as amplified using PCR primer pair Spy-fabM-RF-F and Spy-fabM-RF-R, and the erythromycin resistance gene was amplified using PCR primer pair hsdR-ery-F and hsdR-ery-R using chromosomal DNA of strain NV6 as template (Bjanes et al mBio 2024). PCR products were purified using the DNA Clean and Concentrator Kit (Zymo, Inc.). The three fragments were then ligated together using Golden Gate Assembly with T4 DNA ligase and BsaI-HF2 (New England Biolabs). The ligation was performed in a thermocycler with the following protocol: 1.5 min at 37°; 3 min at 16°; repeated 50X; 5 min at 37°C; 10 min at 80°C; hold at 4°C. The ligation product was gel purified, and further amplified using PCR primer pair Spy-fabM-LF-F and Spy-fabM-RF-R. 1 µg of the purified PCR product was added to 50 µl NV2 competent cells (*S. pyogenes* 5448 containing the pAV488 recombineering plasmid) (Bjånes et al. 2024), which had been thawed on ice. DNA was mixed with the cells and incubated on ice for 5 min, then electroporated at 1600V. 250 µl THB + 0.25 M sucrose was added immediately following electroporation, and the culture was incubated at 37°C for 2 hours and then plated on THB agar + 0.5 µg/ml erythromycin. 8 colonies were restreaked on to THB agar + 0.5 µg/ml erythromycin, and colonies on those plates were used to inoculate 5 ml THB + 0.5 µg/ml erythromycin + 1 mM IPTG to remove the recombineering plasmid. Individual clones from those plates were streaked on to THB agar + 0.5 µg/ml erythromycin and THB agar + 2 µg/ml chloramphenicol to confirm loss of the plasmid. One colony that was positive for erythromycin resistance and negative for chloramphenicol resistance was used to create the frozen stock that was used for all subsequent studies. Desired mutagenesis (i.e., replacement of *fabM* with *ermB*) was confirmed by using the Bacterial Genome Sequencing service by Plasmidsaurus, Inc.

#### S. pneumoniae ΔfabM

A *S. pneumoniae fabM* deletion mutant strain (*SpnΔfabM*) was constructed using a modified method based on a previously described protocol (Hirose et al. 2018). Briefly, Genomic DNA from *S. pneumoniae* TIGR4 (Tettelin et al. 2001) using the Quick-DNA Fungal/Bacterial Miniprep Kit (Zymo, Inc.) according to the manufacturer’s instructions. The upstream and downstream flanking regions of the *fabM* gene were PCR-amplified and fused with an *ermB* resistance cassette derived from the pDCerm plasmid (Jeng et al. 2003) using primers TIGR4-fabM-Up-F, TIGR4-fabM-Up-R, TIGR4-fabM-ermB-F, TIGR4-fabM-ermB-R, TIGR4-fabM-Dw-F, TIGR4-fabM-Dw-R. An isogenic *S. pneumoniae* TIGR4 strain was transformed with the resulting PCR product to generate the *SpnΔfabM* mutant using CSP-mediated natural competence (Bricker and Camilli 1999). Colonies exhibiting erythromycin resistance were identified as *SpnΔfabM,* and proper integration of the *ermB* cassette was confirmed using primers fabM-KO-conf-F and fabM-KO-conf-R and Sanger sequencing.

#### S. pneumoniae fabM-C

A complemented strain carrying a *fabM*-expressing plasmid (*fabM*-C) was constructed using a modified method based on a previously described protocol (Hirose et al. 2018). To construct the *fabM*-expressing plasmid, the *fabM* gene was PCR-amplified with EcoRI and BamHI restriction sites added at the 5′ and 3′ ends, respectively using primers TIGR4-fabM-comp-F, TIGR4-fabM-comp-R, pDC123_universal_F and pDC123_universal_R and ligated into the pDC123 plasmid (Chaffin and Rubens 1998), which carries a chloramphenicol resistance cassette (M. H. Lee et al. 1999). The *SpnΔfabM* strain was transformed with the *fabM*-expressing plasmid using CSP-mediated natural competence (Bricker and Camilli 1999) to generate the complemented strain. Colonies exhibiting resistance to both erythromycin and chloramphenicol were identified as fabM-C.

### S. mutans pcomDE-luc and S. mutans pcomDE-luc-ΔfabM reporter strains

Using overlapping primers pcomDE-R, pcomDE-R, DERluc-F, DERluc-R, T1F5, T1F3, Tuf5, KT13, T1R5, and T1R3, a construct was made using PCR and Gibson Assembly containing 461 bp upstream of *comE* (i.e., the *comCDE* promoter) followed by the open reading frame of Renilla luciferase (*Rluc*), followed by the constitutively active *S. mutans tuf* promoter, followed by a kanamycin resistance gene *kanR,* flanked by 500 bp regions upstream and downstream of the SMU.1180c gene, such that the construct would insert into the SMU.1180c gene. This construct was transformed into *S. mutans* UA159 or *SmuΔfabM* using CSP-induced transformation. Briefly, an overnight culture of *S. mutans* was diluted 1:50 with fresh THB. An aliquot of 200 μL of diluted culture was transferred to a sterile well of 96 well plate and mixed with CSP (500 nM) and target DNA. After incubation at 37C with 5% CO_2_ for another 2h, the culture was streaked on THB plates supplemented with the appropriate antibiotic concentration. Colony formation was assessed after two days.

### S. mutans pnlmA-mCherry. S. mutans pnlmA-mCherry-ΔfabM reporter strains

Using overlapping primers P150-5, P150-3, T2nlmAcherryO5, T2nlmAmcherryO3, N2F5, N2F3, N2R5, and N2R3 a construct was made using PCR and Gibson Assembly containing 212 bp upstream of *nlmA* (i.e., the *nlmA* promoter) followed by the open reading frame of mCherry, followed by the *S. mutans tuf* promoter, followed by a kanamycin resistance gene *kanR,* flanked by 500 bp regions upstream and downstream of the SMU.221c gene, such that the construct would insert into the SMU.221c gene. This construct was transformed into *S. mutans* UA159 or *SmuΔfabM* using CSP-induced transformation. Briefly, an overnight culture of *S. mutans* was diluted 1:50 with fresh THB. An aliquot of 200 μL of diluted culture was transferred to a sterile well of 96 well plate and mixed with CSP (500 nM) and target DNA. After incubation at 37C with 5% CO_2_ for another 2h, the culture was streaked on THB plates supplemented with the appropriate antibiotic concentration. Colony formation was assessed after two days.

## Growth Curves

Overnight cultures of the indicated strains were normalized to the OD_600_ of the lowest density culture in the comparison using the indicated growth media. Growth curves were performed in either 96-well or 384-well clear plates (Corning, Inc.). For growth curves in 96-well plates, 10 µl of the overnight cultures were added to 200 μl of the indicated growth media with or without indicated compounds. For growth curves using 384-well plates, 2.5 µl of the overnight cultures were added to 50 µl of the indicated growth media with or without the indicated compounds. Plates were sealed with Breathe-Easy sealing membrane (Diversified Biotech) and growth curves were performed without the plastic plate lid. Growth was monitored using an Infinite Nano (Tecan Group, Ltd.). Optical density at 600 nm (OD_600_) (and mCherry fluorescence or luciferase luminescence, when applicable) were measured every 30 minutes for the indicated amount of time under 37 °C, with 5 s of shaking prior to each reading. 4-8 replicates of each strain were monitored.

## Derivatization of fatty acid methyl esters for GC-MS

Cultures of the indicated strains and growth media were incubated at 37°C in a 5% (vol/vol) CO_2_–95% air environment for the indicated amount of time. Cells were harvested by centrifugation at 4,000 x g for 10 min. The resulting pellets were decanted and stored at -80° C. Pellets were resuspended and then vortexed in 100 µl concentrated sulfuric acid (96%) and 200 µl methanol. Samples were then boiled by incubation in boiling water for 5 min and allowed to cool to RT. 300 µl of dichloromethane was added, samples were then vortexed and centrifuged 1 min at 15,000 x g. The organic (bottom) layer was transferred to a new tube containing a pinch of Na_2_SO_4_, mixed by vortexing, and centrifuged 1 min at 15,000 x g. The supernatant was then transferred to an HPLC tube and stored at 4°C until being loaded on the GC-MS.

### GC-MS

1 µl of sample was injected into a Shimadzu GC-2010 GC-MS equipped with an AV-1701 capillary GC column (Zebron, Inc.) using an AOC20i auto injector (Shimadzu). The carrier gas was helium and the column oven temperature program was the following: 45°C for 2 min; ramp temperature 9°/min to 180°C then hold for 5 min; ramp temperature 40°/min to 220°C then hold for 5 min; ramp temperature 40°/min to 240°C then hold for 11.5 min; ramp temperature 40°/min to 280°C and hold for 2 min (43 min total). Spectra from 20-650 m/z were collected from 6 min to 37.5 min with a scan speed of 10,000 and an event time of 0.1 s. The ion source temperature was 200°C and the interface temperature was 250°C. .qgd data files were converted to CDF files using GCMS Solutions Postrun Analysis (Shimadzu). CDF files were converted to .mzML files using MZmine (Heuckeroth et al. 2024). Spectral alignment, deconvolution, integration as well as library search was performed using MS-Hub as part of the Global Natural Products Social Molecular Networking (GNPS) GC-MS EI Data Analysis pipeline (Aksenov et al. 2021). Identity of peaks was also confirmed based on comparison of retention times and mass spectra to a known standard component of TraceCERT 37 Component FAME Mix (Supelco).

### Deferred antagonism (i.e., “competition”) assay

The deferred antagonism assay was performed as previously described (Uranga et al. 2021). Briefly, 8 mL of overnight cultures of the initial colonizer was spotted onto THB 1% agar and incubated overnight at 37°C under 5% CO2/95% air. The following day, the plates were sterilized using the sterilization setting (90 s) in a GS Gene Linker UV Chamber (Bio-Rad, Inc.). 500 µL of overnight cultures of *S. sanguinis* SK36, *S. gordonii* ATCC 10558, or *S. mitis* F0392 was added to 5 mL molten THB 0.75% agar that had been cooled to 40°C, and this was used to overlay the plates with the *S. mutans* colonies. The agar overlay was allowed to solidify at room temperature, and then the plates were incubated overnight at 37°C under 5%CO2/95% air. Zones of inhibition were examined and photographed the following day.

### Scanning electron microscopy

Overnight cultures of the indicated strains were used to inoculate 1ml of THB in a 24-well plate (Corning, Inc.) containing a Thermanox disc (Nunc). Plates were incubated overnight at 37°C in a 5% (vol/vol) CO_2_–95% air environment. The following day, media was gently aspirated from the wells, and 1 ml fixing agent (2.5% formaldehyde/glutaraldehyde in 0.1 M sodium cacodylate buffer pH 7.4, [Electron Microscopy Sciences]) was added. Cells were fixed for at least one hour at 4°C. Samples were dehydrated using an ethanol gradient, then a critical point dryer (Leica CPD300) prior to sputter coating with 10 nm-thick carbon (ACE600). Cells were imaged on an Apreo scanning electron microscope (Thermo Fisher Scientific). Sample preparation beyond the fixing, and imaging, were performed at the OHSU Multiscale Microscopy Core, a member of the OHSU University Shared Resource Cores. RRID:SCR_009969.

### Neutrophil killing assay

#### Neutrophil preparation

Blood sample collection protocols were reviewed by the Oregon Health and Science University Institutional Review Board (IRB) prior to the initiation of the study and deemed to be not human subject research (#20759). Venous blood was collected at the Oregon Clinical and Translational Research Institute (OCTRI) from healthy donors and neutrophils were isolated as previously described (Ragland and Criss 2019). Briefly, red blood cells were removed by dextran sedimentation followed by Ficoll gradient centrifugation, and hypotonic lysis. Neutrophils were subsequently resuspended in Phosphate Buffered Saline (without calcium and magnesium) containing 0.1% dextrose and used immediately. <u>Serum preparation.</u> Blood was drawn from a healthy donor as described above and added to a serum tube. The serum tube containing the blood was placed on a rocker for 30 min at RT. The serum was separated from the resulting clot by centrifugation at 1000 x g for 10 min. The serum was then aliquoted into two equal volumes in 15 ml centrifuge tubes. One tube was heat inactivated in a 56° C water bath for 30 min.

#### Neutrophil killing assay

Mid-log phase bacterial cells of the indicated strain were harvested by centrifugation at 4,000 x g for 10 min.

The pellet was decanted and subsequently resuspended in HBSS+/+ buffer such that a an equal 10 µl volume of bacterial cells would have the desired MOI compared to a 10 µl volume of neutrophils. 10 µl of bacteria in HBSS+/+ were added to a round bottom 96 well plate (Corning, Inc.). 5 µl of serum was added to the bacteria and incubated for 30 min on a rocker at RT. 10 µl of neutrophils or 10 µl of HBSS+/+ were then added and the total volume was brought to 100 µl by addition of 75 µl of HBSS+/+. The plate was centrifuged at 500 x g for 5 min. After the indicated amount of time incubating at 37°C, the samples were added to 100 µl of ice-cold 0.3% saponin and incubated on ice for 2 min. The samples were then serially diluted in DMEM out to 1 x 10^-8^, plated on THB agar, and incubated overnight at 37°C in a 5% (vol/vol) CO_2_ –95% air environment.

## Acknowledgements

This research was supported by NIH K99-DE029228 (J.L.B.), R00-DE029228 (J.L.B.), R25-DE032536 (K.C., S.W., M.L., and S.G.D), R25-GM154345 (S.R.), R35-DE028252 (J.M.), and Coordenação de Aperfeiçoamento de Pessoal de Nível Superior – Brasil (CAPES), Finance Code 001 (F.F.F.S.). The authors acknowledge and thank the staff and support of the following Oregon Health & Science University (OHSU) Cores and Institutes: 1) OHSU Multiscale Microscopy Core (a member of the OHSU University Shared Resource Cores; RRID:SCR_009969), especially Raakhee Shankar, for assistance with preparing samples and imaging with scanning electron microscopy, 2) Oregon Clinical and Translational Research Institute (OCTRI; funded by NIH UL1TR002369) for providing the human blood used in this study, and 3) the OHSU Bioanalytical Shared Resource / Pharmokinetics Core, especially Jenny Luo for assistance with the GCMS.

---

**Figure S1:** ***SpnΔfabM* does not produce MUFAs and *Spn-fabM-C* has restored MUFA production.** Panels A-C, Chromatograms from GC-MS of the indicated strain. Peaks are labeled with the mass to charge ratio (m/z; which is approximately 74 for SFAs and 55 for MUFAs), common name, and lipid number. Peaks were identified based on having the same retention time and mass spectra as known standards. Panel D, growth curve illustrating rescue of growth in the *S. pneumoniae fabM-C* (complement) strain.

